# Evaluation of Postgraduate Educational Environment of Doctors Training in Psychiatry: a mixed method study

**DOI:** 10.1101/2022.02.24.481497

**Authors:** Musaab Elzain, Lisa Moran, Geraldine McCarthy, Sarah Hyde, John McFarland

## Abstract

**Introduction:** Educational Environment (EE) is of paramount importance in Medical Education, but can be intangible and hard to clearly determine. Professional satisfaction and patient care improve in a pleasant learning environment; where postgraduate physicians are encouraged, suitably supervised, and fostered. A negative learning environment can be detrimental to trainees’ and teams’ morale and can jeopardize the multidisciplinary working relationship.

**Objectives:** The objective of this study was to measure the Educational Environment (EE) of Psychiatry postgraduate training in the Midwest of Ireland training Deanery; what aspects of training are working well and what areas are seen as not optimal.

**Methods:** This study took place between April and June 2021. A mixed methods approach was adopted, using the Postgraduate Hospital Educational Environment Measure (PHEEM) and semi-structured one-to-one interviews.

**Results:** Response rate was 88% (n=22). The total PHEEM score was 105.64±23.52, indicating a postgraduate EE with more positive than negative aspects, but with room for improvement. There were no differences in overall PHEEM and subscale scores between trainees’ gender, training grades or years of working experience. Three themes were identified that contribute to trainees’ perception of EE: the commitment of the trainees’ supervisors to the role they play in trainees’ overall development, the clinical workload of the trainee, and the day-to-day working conditions of the trainee. Definite disparities between work placements were evident in the collected data across these three themes.

**Conclusions:** Although the training program had an overall positive EE, specific answers and interview themes indicated some areas of weakness that may contribute to trainee dissatisfaction and possible burnout. Planned interventions targeting these areas and tracking changes in EE and burnout rates over time may be useful measures going forward.

**Declaration of Interest:** None

## Introduction

The educational environment(EE) is a metric for evaluating the quality of a curriculum[1] and has been considered the “core” of postgraduate medical education as it encompasses the social, cultural and material context in which trainee doctors learn while working[2, 3]. EE is of paramount importance in Medical Education but is inherently complex and difficult to define in categorical terms [4, 5]. There is evidence that professional satisfaction and patient care improve in a learning environment where postgraduate physicians are encouraged, suitably supervised and fostered[6]. Conversely, a negative learning environment may be detrimental to trainees and team morale, and potentially jeopardizing the multidisciplinary working relationship[7].

The quality of the EE during postgraduate training has consistently been observed to predict the quality of patient care provided by trainees for years after graduation [8, 9]. Furthermore, efforts to enhance the EE have a significant impact on both trainee learning and practice in the future. As trainees represent the frontline of patient care now and the future of health care practice tomorrow, there is a clear rationale for evaluating EEs[8]. Poor EE for training have also been identified as a major driver of physicians’ burnout[7, 10]. There is consistent evidence for a link between burnout and the learning environment for postgraduate medical training, with negatively perceived learning environments associated with increased burnout[11, 10].

### Psychiatry Training in Ireland

Psychiatry training in Ireland has undergone significant change over recent years, with many developments positive for trainees. Irish psychiatry postgraduate training follows a Competency Based Learning curriculum[12]. The College of Psychiatrists of Ireland (CPsychI) distinguishes between clinical and educational supervision. Clinical supervision promotes trainee growth while assuring the best possible patient care and it must be offered at a level that is appropriate for the trainee’s needs and experience. Educational supervision, has a distinct, separate focus on career and training planning, mentoring, coaching and ensuring that learning objectives are met and documented[12].

To our knowledge, no study has looked into the EE of psychiatry training in Ireland. The current study sets out to gain an understanding of how the psychiatry trainees in the Midwest of Ireland perceive their educational environment.

## Study aims and objectives

### Objective

To measure the EE of psychiatry postgraduate training in the Midwest of Ireland training Deanery.

### Aims

a. To evaluate trainee perceptions of their EE; what aspects of training are working well and what areas are seen as not optimal.
b. To compare perceptions among trainees of different grade, gender and levels of working experience.

## Methods

### Study design

The study was conducted in the Psychiatry Department at the University of Limerick between April and June 2021. A mixed-methods approach was taken with quantitative and qualitative data collected and analysed sequentially. The use of a mixed methods approach allowed for more thorough and meaningful answers to the research question. The Postgraduate Hospital Educational Environment Measure (PHEEM)[13] was developed to assess postgraduate EE and has been administered to many groups, including trainees at varying levels and in different specialties including psychiatry. It has shown to be valid, reliable and practical and is the de facto standard [14,6,15]. Therefore, the PHEEM was deemed the most suitable measurement tool for this study.

The University of Limerick Psychiatric Training Scheme/ Deanery (1 of 9 deaneries in Ireland) is responsible for the training of approximately 25 psychiatry trainees. The Trainees are assigned to training approved clinical posts in various psychiatry specialities across the Midwest of Ireland. The region serves a population of 470,000 people, who live in three counties in a mix of urban and rural settings. Following ethical approval, an anonymised PHEEM survey was sent to all eligible participants (n= 25) in April 2021 with a thirty-day response window.

### The PHEEM Questionnaire

The PHEEM consists of 40 items divided into three subscales: perceptions of role autonomy, perceptions of teaching, and perceptions of social support [13]. Participants were asked to think carefully about each statement before responding on a 5-point Likert scale ranging from “strongly agree” to “strongly disagree”. The PHEEM questionnaire’s authors proposed the following guidelines for evaluating the total PHEEM score: (0-40) Very poor, (41–80) Plenty of problems, (81–120) More positive than negative but room for improvement and (121–160) Excellent. Table 1 shows the PHEEM items belonging to each sub-scale.

**Table 1.** Responses to individual PHEEM items.

| *Item* | *Strongly agree (%)* | *Agree (%)* | *Uncertain (%)* | *Disagree (%)* | *Strongly disagree (%)* | *Mean±SD* |
| --- | --- | --- | --- | --- | --- | --- |
| I have a contract of employment that provides information about hours of work | 50 | 50 | 0 | 0 | 0 | 3.50±0.51 |
| My clinical teachers set clear expectations | 18.2 | 36.4 | 45.5 | 0 | 0 | 2.73±0.78 |
| I have protected educational time in this post | 45.5 | 36.4 | 13.6 | 0 | 4.5 | 3.18±1.00 |
| I had an informative induction programme | 9.1 | 40.9 | 18.2 | 18.2 | 13.6 | 2.14±1.25 |
| I have been given an appropriate level of responsibility in this post | 27.3 | 68.2 | 0 | 4.5 | 0 | 3.18±0.66 |
| I have good clinical supervision | 10 | 31.8 | 13.6 | 4.5 | 4.5 | 3.09±1.11 |
| *There is racism* | 27.3 | 0 | 45.5 | 27.3 | 0 | 2.82±0.85 |
| *I have to perform inappropriate tasks* | 4.5 | 13.6 | 18.2 | 45.5 | 18.2 | 2.59±1.10 |
| There is an informative Junior doctor's handbook | 4.3 | 27.3 | 36.4 | 27.3 | 4.5 | 2.00±0.98 |
| My clinical teachers have good communication skills | 27.3 | 54.5 | 18.2 | 0 | 0 | 3.09±0.68 |
| *I receive phone calls inappropriately* | 18.2 | 31.8 | 13.6 | 27.3 | 9.1 | 1.77±1.31 |
| I am able to participate actively in educational events | 13 | 69.6 | 13.6 | 0 | 0 | 3.00±0.54 |
| *There is sex discrimination in this post* | 0 | 0 | 22.7 | 45.5 | 31.8 | 3.09±0.75 |
| There are clear clinical protocols | 9.1 | 36.4 | 40.9 | 13.6 | 0 | 2.41±0.85 |
| My clinical teachers are enthusiastic | 27.3 | 45.5 | 9.1 | 9.1 | 9.1 | 2.73±1.24 |
| I have good collaboration with other doctors in my grade | 18.2 | 45.5 | 9.1 | 13.6 | 13.6 | 2.41±1.33 |
| My hours conform to the European Working Time Directive | 27.3 | 68.2 | 4.5 | 0 | 0 | 3.23±0.53 |
| I have the opportunity to provide continuity of care | 18.2 | 54.5 | 18.2 | 9.1 | 0 | 2.82±0.85 |
| I have suitable access to career advice | 13.6 | 36.4 | 27.3 | 22.7 | 0 | 2.41±1.01 |
| This hospital has good quality accommodation when on call | 13.6 | 9.1 | 22.7 | 18.2 | 36.4 | 1.45±1.44 |
| There is access to an educational programme relevant to my needs | 13.6 | 36.4 | 22.7 | 22.7 | 4.5 | 2.32±1.13 |
| I get regular feedback from seniors | 13.6 | 45.5 | 13.6 | 18.2 | 9.1 | 2.36±1.22 |
| My clinical teachers are well-organised | 22.7 | 40.9 | 18.2 | 9.1 | 9.1 | 2.59±1.22 |
| I feel physically safe within the hospital environment | 18.2 | 68.2 | 4.5 | 9.1 | 0 | 2.95±0.79 |
| There is a no-blame culture in this post | 4.5 | 36.4 | 40.9 | 9.1 | 9.1 | 2.18±1.01 |
| There are adequate catering facilities when I am on call | 4.5 | 13.6 | 27.3 | 27.3 | 27.3 | 1.41±1.18 |
| I have enough clinical learning opportunities for my needs | 22.7 | 45.5 | 4.5 | 22.7 | 4.5 | 2.59±1.22 |
| My clinical teachers have good teaching skills | 27.3 | 31.8 | 13.6 | 22.7 | 4.5 | 2.55±1.26 |
| I feel part of a team working here | 31.8 | 45.5 | 4.5 | 13.6 | 4.5 | 2.86±1.17 |
| I have opportunities to acquire the appropriate practical procedures for my grade | 27.3 | 40.9 | 31.8 | 0 | 0 | 2.95±0.79 |
| My clinical teachers are accessible | 36.4 | 50.0 | 13.6 | 0 | 0 | 3.23±0.69 |
| My workload in this job is fine | 13.6 | 54.5 | 9.1 | 18.2 | 4.5 | 2.55±1.10 |
| Senior staff utilise learning opportunities effectively | 13.6 | 54.5 | 18.2 | 13.6 | 0 | 2.68±0.89 |
| The training in this post makes me feel ready to be a SPR/Consultant | 22.7 | 31.8 | 31.8 | 13.6 | 0 | 2.64±1.00 |
| My clinical teachers have good mentoring skills | 18.2 | 50.0 | 13.6 | 18.2 | 0 | 2.68±1.00 |
| I get a lot of enjoyment out of my present job | 13.6 | 54.5 | 22.7 | 9.1 | 0 | 2.73±0.83 |
| My clinical teachers encourage me to be an independent learner | 18.2 | 77.3 | 4.5 | 0 | 0 | 3.14±0.47 |
| There are good counselling opportunities for junior doctors who fail to complete their training satisfactory | 0 | 4.5 | 68.2 | 18.2 | 9.1 | 1.68±0.72 |
| The clinical teachers provide me with good feedback on my strengths and weaknesses | 22.7 | 54.5 | 4.5 | 13.6 | 4.5 | 2.77±1.11 |
| My clinical teachers promote an atmosphere of mutual respect | 31.8 | 54.5 | 9.1 | 4.5 | 0 | 3.14±0.77 |

Minor modifications were made to the PHEEM to reflect subtle differences between the UK and Irish context; item 17 ‘*my hours conform to the new deal*’ was replaced with ‘*my hours conform to the European Working Time Directive*’, and item 11 ‘*bleeped*’ was replaced with ‘*phone calls*’. Demographic information on gender, training grade, number of years in current grade was collected.

### Statistical Analysis

Statistical analysis was carried out using the Statistical Package for Social Sciences 26.0 for Windows (SPSS, IBM, and Armonk, NY, USA). Data was presented as mean and standard deviation for quantitative data, and proportions for categorical data. Unpaired Student’s t-test was used for parametric quantitative data comparisons and Wilcoxon signed-rank test for nonparametric quantitative data. Differences were considered significant if p < 0.05. Reliability analysis was performed using Cronbach’s alpha coefficient.

### Qualitative phase, one-to-one individual interviews

Semi-structured one-to-one interviews were conducted to supplement the PHEEM results. All trainees (n = 25) were invited by e-mail to participate in interviews, purposeful sampling of eight psychiatry trainees were selected to ensure heterogeneity. The selected trainees were working in different psychiatry sub-specialities and/or different stages of training. Consent was obtained from the participants for audio recording and verbatim transcribing of the interviews. Questions were designed to gain understanding of the trainee’s personal experience of the EE and to understand factors involved in trainee’s positive and negative experiences. An iterative approach was used, with questions focusing on areas that arose from the PHEEM and interviews. Interviews lasted from 45 - 75 minutes and continued until data saturation was reached (no new themes were identified from the final three interviews). Interviews were conducted and transcribed by the first author (MZ). Interviewees were given a summary of the interviews to confirm accuracy.

Data familiarisation was achieved by listening to the audiotapes and reading the full data sets several times. Two authors (MZ and LM) used an open coding strategy to reduce the data and uncover the basic concepts. Similar concepts were grouped together to form themes. Differences in concepts were resolved through discussion between the authors. An inductive approach was adopted, using data from the one-to-one interviews to triangulate concepts and draw conclusions. The research team documented the entire research process, including the development of the potential participants’ database, the interaction with these participants for data collection, the data analyses, and the engagement among the different co-authors, to ensure transparency and develop an audit trail of research developments.

The primary researcher is a post-graduate trainee in the final year of the Higher Speciality Training (HST). He completed four-year Basic Speciality Training (BST) and subsequently enrolled in the HST at University of Limerick Training Scheme. As a result, he has a strong awareness of the national and local psychiatry training landscape, while holding minimal power over any of the postgraduate trainees. It was anticipated that trainees would feel comfortable discussing their experiences with him and this relationship was further validated by regular reflexive use of memos, as recommended by Charmaz[16], to understand and minimize potential for unintended impacts upon data analysis. The remainder of the research team includes four experienced researchers holding advanced degrees (MSc, MD), from diverse backgrounds with different theoretical perspectives (Psychiatry, General Practice and Medical Education). The team contributed to the research protocol development, data analysis and methodological rigour on an ongoing basis.

## Results

Response rate for completed PHEEM questionnaires was 88% (n = 22). 50% of respondents were female. Cronbach’s alpha was 0.958, and when analysed to exclude each question, no significant improvement in the score was obtained, reflecting high internal reliability and no irrelevant questions.

### Characteristics of participants

Of the 22 respondents 77 %(n=17) were BST and 23 %(n=5) were HST. 41 %(n =9) of respondents had worked for 1 year in their current training grade, 22 %(n = 5) and 36 %(n=8) trainees had worked for two and three years in their current training grade, respectively.

### The PHEEM Score

Table 1 summarises the mean responses to each item on the PHEEM questionnaire. The total PHEEM score was 105.64±23.52 (Range=56-155), corresponding to an EE with more positive than negative aspects, with room for improvement. Findings were comparable with other educational institutions’ PHEEM scores in recent years (table 2). No trainee scored the EE as very poor. 71% (n=12) of BST and 60 %(n=3) HST trainees scored the EE as more positive than negative. 12 %(n=2) of BST trainees and one HST trainee scored it as having significant problems. 12 %(n=2) of BST trainees and one HST trainee scored the EE as excellent.

**Table 2.** PHEEM comparable with other educational institutions PHEEM scores in recent years.

| ***Reference*** | ***Country*** | ***Speciality*** | ***Participants Number*** | ***Total PHEEM*** | ***Autonomy*** | ***Teaching*** | ***Social support*** |
| --- | --- | --- | --- | --- | --- | --- | --- |
| *Elzain , 2022* | Ireland | Psychiatry | 22 (88%) | 105.0 | 37.7 | 42 | 25.8 |
| Ong , 2020[17] | Singapore | Internal Medicine | 136 (88.9%) | 112.23 | 38.5 | 42.79 | 30.93 |
| Gough , 2010[18] | Australia | Junior doctors | 429 | 110.0 | NA | NA | NA |
| Mahendran, 2013[19] | Singapore | Psychiatry | 60 (78.9%) | 109.3 | NA | NA | NA |
| Clapham , 2007[20] | UK | Intensive Care | 134(71%) | 103.5 | 35.7 | 38.3 | 28.4 |
| Buali, 2015[21] | KSA | Paediatrics | 104 (86.7%) | 100.19 | 34.91 | 38.89 | 26.38 |
| Flaherty, 2016[6] | Ireland | Internal Medicine | 61 (60%) | 82.9 | NA | NA | NA |
| T. Khoja1 , 2015[22] | KSA | Family Medicine | 101(91.1%) | 67.1 | 24.2 | 1.9 | 28.4 |
| Shimizu, 2013[23] | Japan | Internal Medicine | 206 | 57.6 | NA | NA | NA |

We found no statistically significant differences in overall PHEEM and subscale scores between genders and trainees’ grades. The lowest-scoring items were: Question 26, ‘There are adequate catering facilities when I am on call’ (mean 1.41±1.18); Question 20, ‘This hospital has good quality accommodation when on call’ (1.45±1.44); Question 38, ‘There are good counselling opportunities for junior doctors who fail to complete their training satisfactory’ (mean 1.68±0.72) and Question 11, ‘I receive phone calls inappropriately’ (mean 1.77±1.3).

### PHEEM subscale scores

Perception of role autonomy PHEEM score was 37.77 (Range 23-54) corresponding to score of a more positive perception of one’s job, perception of teaching PHEEM score of 42.04 (Range 21-60) corresponding to score of moving in the right direction, and perception of social support PHEEM score of 25.8 (Range 12-42) corresponding to score of more pros than cons (table 2).

### Qualitative part, one-to-one individual interviews

The participants were one Year 1 BST, two Year 2 BST, two Year 3 BST, one Year 4 BST and two HST. Four (50%) were female. Three themes were identified that contribute to trainees’ perception of EE: the commitment of the trainees’ supervisors to the role they play in trainees overall development, the clinical workload of the trainee, and the day-to-day working conditions of the trainee.

1. The commitment of the trainees’ supervisors to the role they play in trainees overall development

### Trainee-supervisor relationship

Interactions between supervisors and trainees were discussed by all trainees and identified as a central factor for trainees’ development. Relationships with other team members was also discussed, but the trainee-supervisor relationship was the most impactful (Table 3). Most trainees described a positive relationship with their supervisors, but a poor/negatively described relationship made their job more difficult and less enjoyable.

**Table 3.**
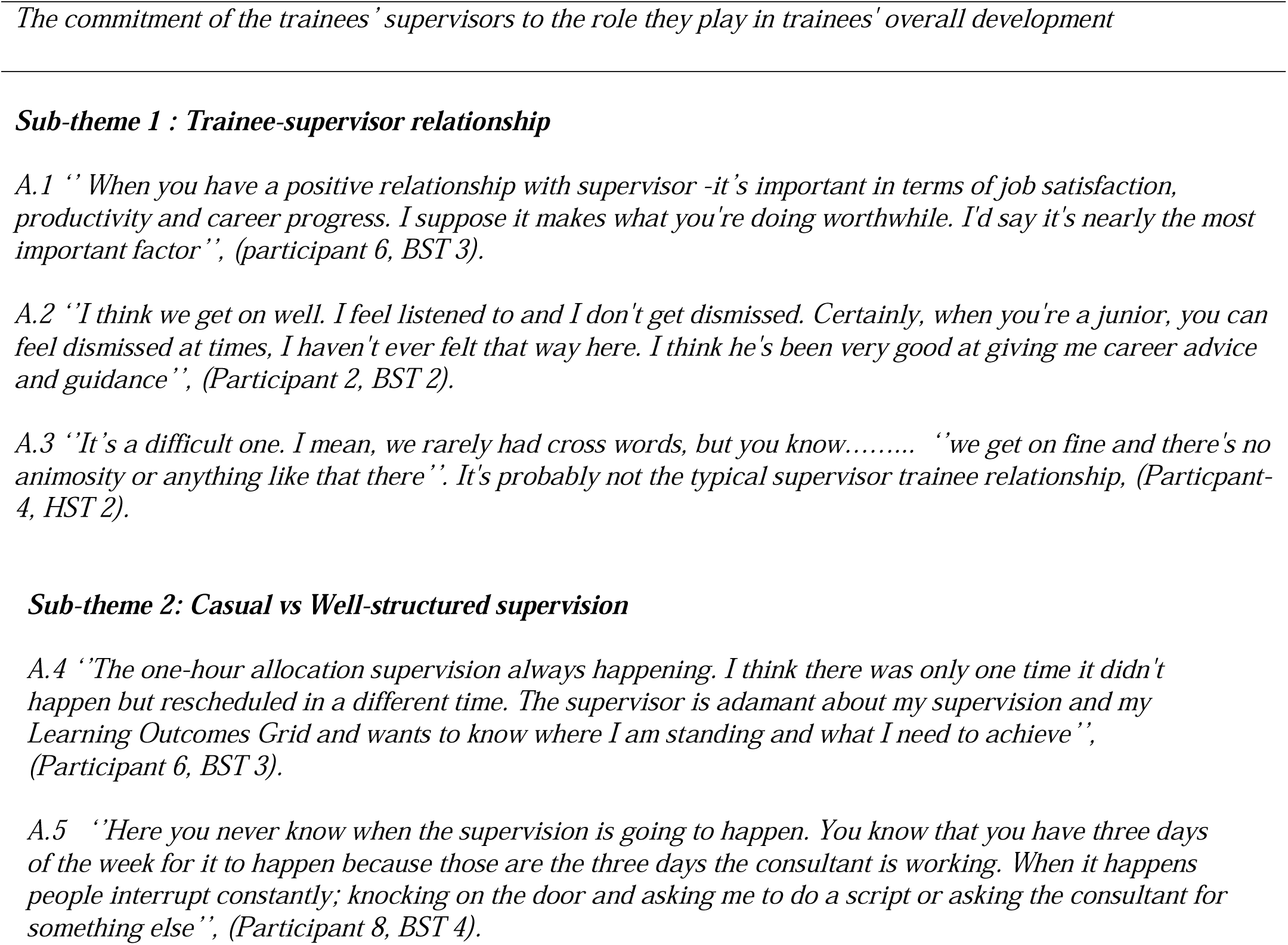
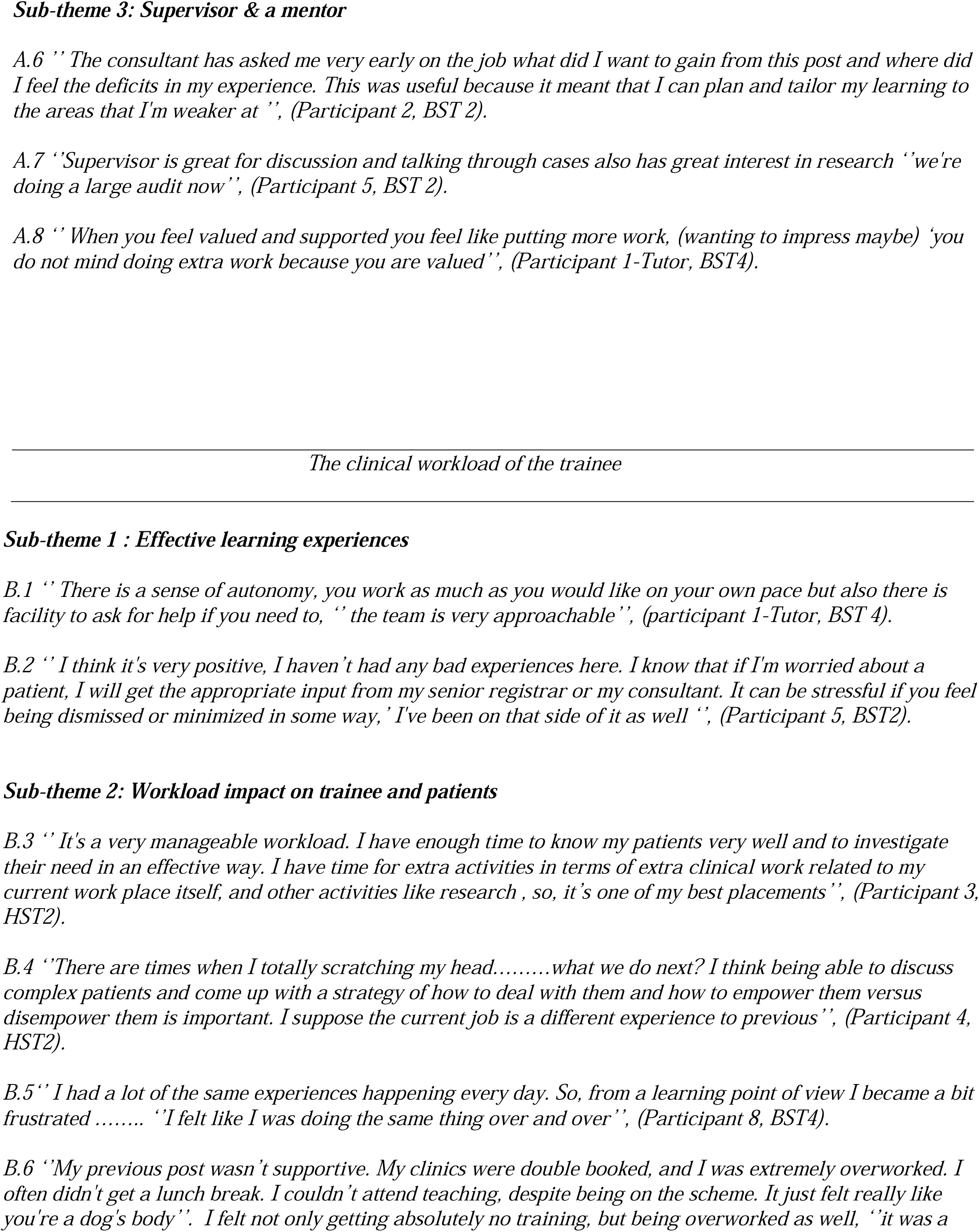
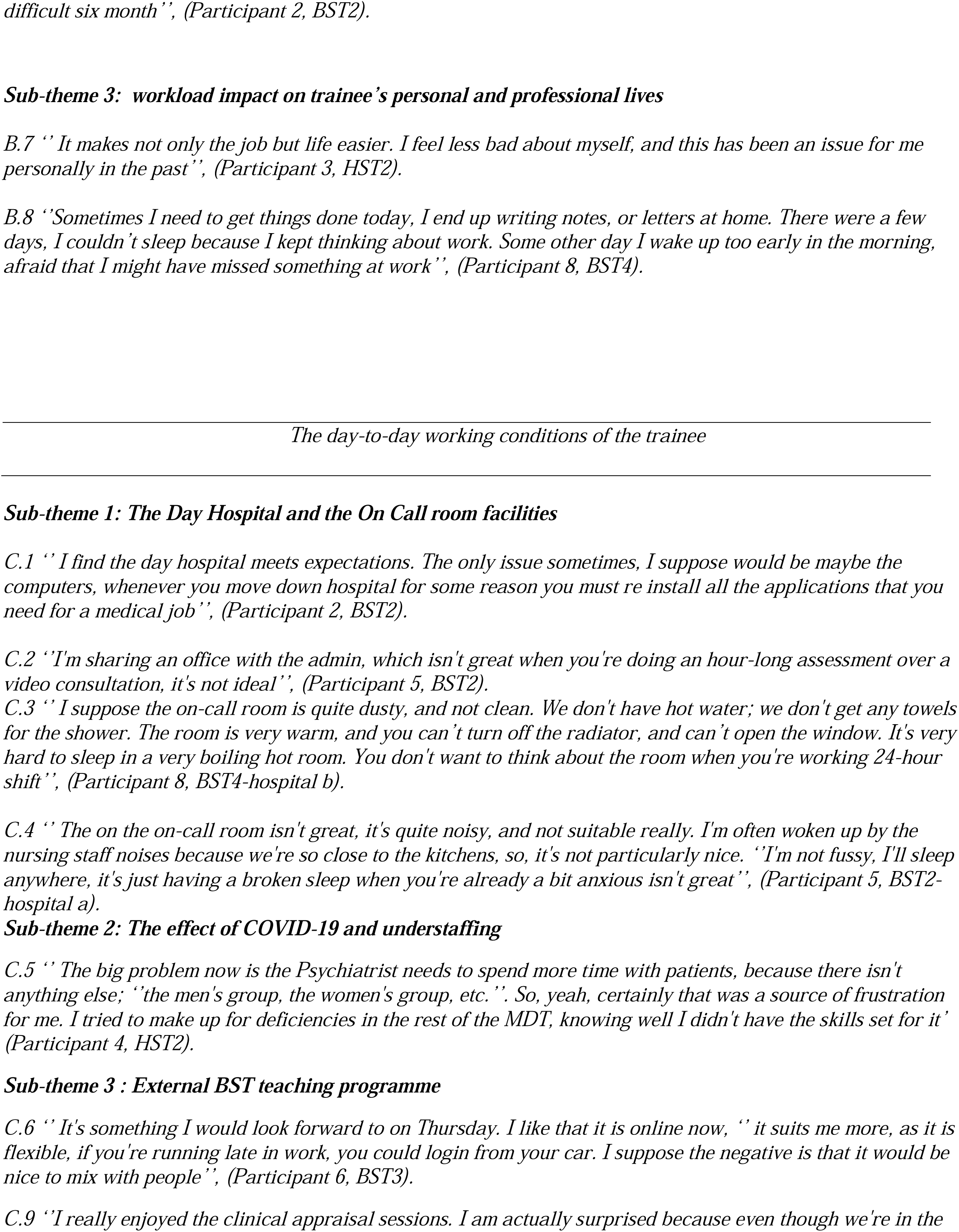
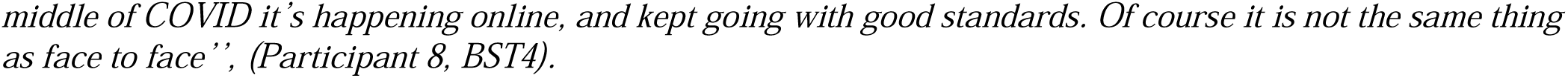
Quotes relating to the theme 1, 2& 3.

Supervisors being supportive and valuing a trainee’s education and development led to greater job satisfaction among trainees. The availability of supervisors for advice on clinical questions helped trainees learn and feel supported making decisions. This was in contrast to when supervisors did not explain their clinical reasoning, or share their thought processes, e.g. when prescribing new medications.

### Casual vs Well-structured supervision

Trainees were appreciative when supervisors took their supervision role seriously, making them feel that their education was important. Conversely, some supervisors took a more casual approach to supervision, which impacted on trainees’ impression of their role.

### Supervisor and a mentor

Trainees described some supervisors taking on a mentor role, providing career guidance and research opportunities. Some supervisors were encouraging of trainees to tailor their learning to their specific learning needs. When trainees felt supported and valued in their role on the team, they felt inclined to go above and beyond their stated duties in order to impress their supervisors.

2. The clinical workload of the trainee

### Effective learning experiences

Trainees who had an interesting, varied clinical workload with a supportive team enjoyed their job and felt that their learning was preparing them for their future roles. Effective clinical learning experiences included: supervisors being supportive after an adverse outcome, supervisors respecting the trainee’s clinical opinion, and when trainees were given autonomy to make decisions. This was felt to accelerate learning (Table 3).

### Workload impact on trainee and patients

Trainees reported that their clinical workload and sense of support could influence patient care, either in a positive or a negative way. Trainees experiencing a supportive atmosphere felt more secure when dealing with challenging patients, whereas those in a less supportive environment felt befuddled and unsure of what to do. It was apparent that Trainees value broad exposure to different presentations in psychiatry, and the opportunity to work with others in order to learn. Those with a less varied clinical workload expressed frustration at times. Some trainees felt that their current post prepared them for the future and shaped their career, however others described clinical workloads that were excessive and impacted negatively on their learning.

### Workload impact on trainee’s personal and professional lives

The impact of a manageable and challenging workload with a supportive team was felt by trainees beyond their working day and into their personal and professional lives. Conversely, trainees who were overworked and unsupported described significant impacts on their personal lives.

3. The day-to-day working conditions of the trainee

*All the trainees reported that their weekly working hours adhered to the European Working Time Directive (EWTD)*.

### The Day Hospital and the On Call room facilities

*Almost all the trainees described the Day Hospital facilities, where they have their clinics as suitable and appropriate. One trainee mentioned that they shared their office with a member of administrative staff. The On Call room was described by all of the trainees as suboptimal and insufficient. It was difficult to get basic necessities like hot water and towels for a shower. Due to the lack of domestic cleaning on weekends, trainees worried they could encounter COVID-19 (Table 3)*.

### The effect of COVID-19 and understaffing

*Due to understaffing or the COVID-19 effect, some trainees felt they have been placed in situations where they have had to fulfil other team duties*.

### The BST teaching

*External BST teaching is a fortnightly extra-curricular programme facilitated by Consultants and Senior Registrars. BST trainees expressed satisfaction with the programme, despite the changes necessitated by Covid-19*.

## Discussion

### Summary of Findings

This study measured how psychiatry postgraduate trainees perceive their educational environment in one Irish healthcare region. The response rate of 88% to the PHEEM may indicate that trainees were motivated to reflect on and improve their educational environment. In addition to determining how trainees perceive an institution’s EE, the literature suggests that determining what type of EE trainees desire may be of significance[4,24,25]. Till [25] proposes that EE studies that examine the disparity between *real* and *ideal* EE may be utilized to develop short- and long-term strategic approaches for improving healthcare professionals’ educational experiences.

Compared to the published results of other training programmes worldwide, this PHEEM score is overall high, particularly with respect to the perception of autonomy and perception of teaching subscales (Table 2). The College of Psychiatrists of Ireland [12] places emphasis on both clinical and educational supervision and it is possible that this has led to a relatively high level of trainee satisfaction with these areas as seen in the PHEEM. It is notable that this study scored less than other programmes in terms of social support.

One of the disappointing but perhaps unsurprising findings was that three of the lowest four-scoring items were within the social support subscale; Q26, ‘There are adequate catering facilities when I am on call’, Q20, ‘This hospital has good quality accommodation when on call’, and Question 38, ‘There are good counselling opportunities for junior doctors who fail to complete their training satisfactory’. Trainees also scored low on Q11, ‘I receive phone calls inappropriately ‘. Poor on-call facilities and inappropriate interruptions were also discussed by trainees during the interviews. This is an important (and easily addressable) observation, with previous analysis demonstrating that poor working conditions are a major factor in medical emigration and that the possibility of health professionals returning is strongly reliant on improvements in working conditions [26].

27% of the participants in this study felt there is racism in their current post. 45 % of the participants responded uncertain to this question. It is notable that these highly concerning findings were not raised at the qualitative interviews. Nonetheless, trainees may have found it difficult to address these issues in person. In a survey of 476 UK trained doctors, 62 % of doctors from an ethnic minority believed ethnicity had implications for medical training, 70% for early career opportunities, 87% for access to specialties and 86% for career advancement[27].

Our qualitative analysis identified three recurring themes contributing to trainee perception of the EE: the commitment of the Trainees’ supervisors to the role they play in Trainees’ overall development, the clinical workload of the trainee, and the day-to-day working conditions of the trainee. These themes might be used to form the basis of interventions to improve the EE in the training programme. Definite disparities between work placements were evident in the collected data across these three themes.

Trainees who worked in a positive environment reported feeling less stressed in their personal and professional life. Trainees in a less supportive atmosphere reported considerable disruption on personal life and showed warning signs of burnout. In their recent meta-analysis, Panagioti et al. [28] concluded that burnout is an issue for the entire healthcare organization, rather than individuals’ traits and our study provides further evidence for the role of a direct organisational approach in reducing the condition.

Many trainees in this study described very beneficial supervision. The CPsychI places an emphasis on weekly supervision sessions[12], however, the content is dictated by the supervisor or the trainee, with the potential of limited structure. It was notable that half of the trainees in our study described the one hour allocated supervision as casual and lacking clear direction. This perceived lack of clarity for is in line with previous studies from the UK, suggesting that clinical supervisors may not be adequately trained in supervision[29, 30] and that the weekly timetable is not adhered to consistently[31].

Nonetheless, trainees placed a high value on support from supervisor and senior colleagues; in terms of supervision, giving advice and they appeared to particularly value when a supervisor takes a mentor role. Indeed, one of the most important aspects that influenced participants’ perceptions of the EE was the relationship with their supervisor, a finding that is consistent with previous research[25,32,18]. Wasson et al.[33]in a systematic review found that faculty mentor programs were highly rated by students as a technique of preventing burnout. Our study provides further support for programs that emphasize the development of pedagogical competences for providing feedback and effective supervision, as well as mentoring[32].

Autonomy was another recurring theme with increased sense of autonomy resulting in increased trainee satisfaction with EE. Sawatsky et al. [34] explored the tension between autonomy and supervision through the lens of Social Cognitive Theory. They emphasized, in order to create the best learning environments for doctors’ professional identity formation, educators must strike a balance between autonomy, supervision and patient safety.

This developing sense of identity may have been a contributing factor to the expressed satisfaction with the external BST teaching programme. In the Mid-West Deanery, this fortnightly programme has consciously steered away from core Learning Objectives and places an emphasis on areas of reflection, complexities of psychiatry and professional identity. This group has evolved over time, reflecting a Community of Practice (CoP) philosophy [35], where learning involves a process of socialisation in which newcomers to a group move from peripheral to full participation, eventually becoming part of the shared practices and beliefs common to that group process. This evolution may have played a role in the high levels of engagement with the current study, in that participants felt empowered to be agents of change within the Deanery.

Interestingly, the shift of the teaching to an on-line platform was viewed positively by a number of trainees and may have led to improved attendance and participation; a finding that is at odds with the evolving literature base on the COVID-19 pandemic’s impact on medical education [36]. Nonetheless, due to the short time window and unexpected nature of the pandemic, our study provides a limited contribution to the evolving literature on the subject, which largely consists of anecdotal communications with little empirical evidence. To determine the effect of the pandemic in the EE, more research is required with a need for qualitative and quantitative studies.

### Strength and Limitations of the present study

The strength of this study was using a mixed methods analysis, which made our findings more robust with more confidence in our conclusions. The in-depth interviews allowed us to explore the EE; difficult to define concepts that are not easily explained by quantitative data. Our response rate (88%) was high, but small sample is a limitation. This study only investigated a training programme within a single health region in Ireland, which may limit the generalisability of our data. A single researcher who was a trainee member conducted the interviews, however, this effect was reduced by the primary researchers engaging regular reflexive memos.

## Conclusion

Although our training program had a positive EE, specific questions and interview themes indicated some areas of weakness that may have contributed to trainees’ dissatisfaction and possible burnout. Planned interventions targeting these areas and tracking changes in EE and burnout rates over time may be useful measures. There is scope for further qualitative research to explore how the supervisors/trainers perceive the EE to explore the interplay between trainees’ individual and organizational factors. This may help to identify critical periods for intervention and inform the development of a curriculum that promotes resilience and provides targeted supports to prevent burnout. There is an onus on educators and healthcare managers to work towards providing an environment that balances the demands of service delivery with the educational needs of trainees to support their professional development.

## Acknowledgement

The authors wish to acknowledge the generosity of Mrs. Sue Roff and Dr. Sea’n McAleer from the Centre for Medical Education at Dundee University in relation to use of the PHEEM instrument, and the psychiatry trainees at University Hospital Limerick who participated in the study.

## Ethical Standards

The authors assert that all procedures contributing to this work comply with the ethical standards of the relevant national and institutional committee on human experimentation with the Helsinki Declaration of 1975, as revised in 2008. The study protocol was approved by the Ethics Committee of University Hospital Limerick.

Written informed consent was obtained from all the participants.

